# MitoSort: robust demultiplexing of pooled single-cell genomics data using endogenous germline mitochondrial variants

**DOI:** 10.1101/2023.04.26.538392

**Authors:** Zhongjie Tang, Weixing Zhang, Peiyu Shi, Sijun Li, Xinhui Li, Yicong Xu, Yaqing Shu, Jin Xu

## Abstract

Multiplexing across donors has emerged as a popular strategy to increase throughput, reduce costs, overcome technical batch effects, and improve doublet detection in single-cell genomic studies. Using endogenous genetic barcodes eliminates the need for additional experimental processing steps. Among the available choices for endogenous barcodes, the unique features of mtDNA variants render them a more computationally efficient and robust option compared to genome variants. Here we present MitoSort, a method that uses mtDNA germline variants to assign cells to their donor of origin and identify cross-genotype doublets. We evaluated the performance of MitoSort by *in silico* pooled mtscATAC-seq libraries and experimentally multiplexed data using cell hashing method. MitoSort achieve both high accuracy and efficiency on genotype clustering and doublet detection for mtscATAC-seq data, which fills a void left by the inadequacies of current computational techniques tailored for scRNA-seq data. Moreover, MitoSort exhibits versatility and can be applied to various single-cell sequencing approaches beyond mtscATAC-seq, as long as the mtDNA variants can be reliably detected. Furthermore, through a case study, we demonstrated that demultiplexing 8 individuals assayed at the same time with MitoSort, enables the comparison of cell composition without batch effects.

## INTRODUCTION

The advent of single-cell sequencing techniques has expanded the repertoire of detectable modalities within a single cell (1, 2). For instance, DOGMA-seq (3) enables the simultaneous measurement of gene expression, chromatin accessibility, and protein levels in individual cells, providing a powerful tool for the comprehensive exploration of biological processes. However, the high costs of multimodal single-cell techniques limit their applications in large cohort studies. To meet the growing demands for profiling large-scale samples, multiplexing samples has become a popular strategy that not only reduces costs but also overcomes technical batch effects and improves doublet detection (4). To avoid additional experimental processing steps, several computational tools have been proposed to harness genome variants for demultiplexing cells from different individuals and identifying cross-genotype doublets in scRNA-seq data (5–9). This process typically requires significant computational resources, even with reference genotypes, due to the large size of the human genome. Furthermore, the performance of these methods may be suboptimal when applied to single-cell sequencing approaches targeting genome DNAs, such as single-cell ATAC-seq, as their efficacy relies on the read coverage of genome variants.

In contrast to the genome variants, mtDNA variants offer a more computationally efficient and robust option, given their hundreds of copies per cell and the small size of the mtDNA genome (10, 11). Moreover, mtDNA variants can be captured with high coverage at single-cell resolution in multimodal single-cell sequencing techniques such as mtscATAC-seq (12), ASAP-seq (3), DOGMA-seq (3), as well as full-length single-cell RNA-seq (e.g., Smart-seq (13)).

Despite the benefits of using mtDNA variants for demultiplexing samples and identifying doublets, few software tools are currently available. To fill the gap, here we developed MitoSort, a software that exploits mtDNA germline variants exclusively to demultiplex samples from different individuals and detect cross-genotype doublets efficiently and accurately (Figure 1a).

**Figure 1.**
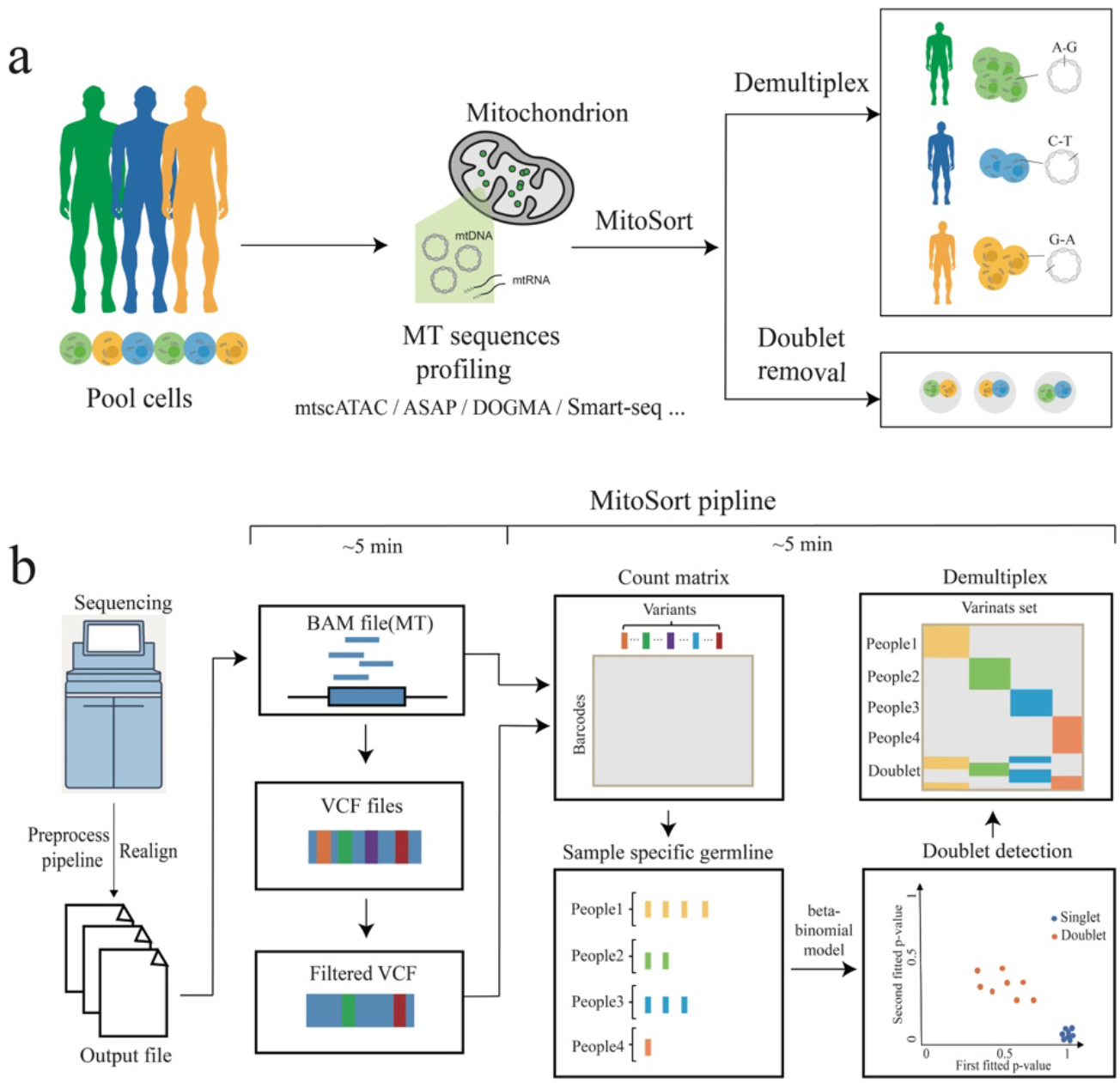
Workflow of MitoSort. **a**, Schematic showing the purpose of MitoSort. **b,** Overview of MitoSort pipeline. Firstly, mitochondrial reads are realigned and putative single-nucleotide polymorphisms (SNPs) are called. Subsequently, count matrices of reference and alternate alleles are generated for each cell, where each row represents a cell and each column represents a variant. Following that, sample-specific germline variants are identified for each individual. Cross-genotype doublets are then detected by a beta-binomial model. Lastly, singlets are demultiplexed for downstream analyses.

We have evaluated the performance of MitoSort using *in silico* pooled and cell hashed mtscATAC-seq data, which demonstrate that MitoSort is a highly accurate and scalable method. MitoSort can be applied to various single-cell sequencing approaches beyond mtscATAC-seq, provided that the mitochondrial sequence can be readout. Besides, the application of MitoSort to a case study further demonstrates MitoSort can be used to accurately demultiplex samples of individuals under different pathologies, which facilitates downstream analyses without technical batch effects.

## MATERIAL AND METHODS

### Realignment for Mitochondrial reads

To correct potential mapping errors around indels, the reads mapped to mitochondrial genome were realigned locally using GATK version 3.5 (14).

### Variant calling and germline mutation selection

VarScan2 (15) was employed with the parameters “–min-var-freq 0.01, min-depth 8 and –min-reads2 2” to call variants on the mitochondrial genome using BAM files containing read pairs realigned to the mitochondrial genome. Since mitochondrial reads from the BAM file were contributed by multiple individuals, we retained variants with allele frequencies ranging from 1% to 99% as potential germline variants for each individual (1% < bulk VAF < 99%). To determine the number of reference alleles and alternate alleles in potential germline variants for each cell, we developed an in-house Python script (available at https://github.com/tangzhj/MitoSort) to generate two matrices, namely, Ref.matrix and Alt.matrix. For each potential germline variant per cell, the allele frequency was calculated as the ratio of alternate counts to all counts and stored in the Frequency.matrix. The Frequency.matrix contained the allele frequency for each potential germline variant, including sample-specific germline variants, common germline variants, and widely spread somatic variants.

To ensure that only germline variants were included in our analysis, a straightforward filtering criterion was employed. Specifically, we selected variants where the proportion of cells having a variant allele frequency (VAF) above 99% among those with a non-zero VAF was higher than 50%. This criterion was chosen to ensure the selection of high-confidence germline variants with a high VAF in most cells, while simultaneously excluding low-coverage germline variants and somatic variants with low VAF.

### Identification of specific germline variants for each individual

To identify sample-specific germline variants while excluding common germline variants that could potentially interfere with subsequent doublet detection, we utilized the robust information on germline variants in each sample. First, we performed initial clustering using the k-means algorithm from the sklearn library, with parameters n_clusters = pooled_sample_number, max_iter=1000, tol=1e-5, algorithm=’full’, and n_init = 10, based on the information obtained from the Frequency.matrix. Within each cell cluster, variants where the ratio between cells with VAF>99% and cells with VAF>0 was higher than 0.5 were assigned as a variant set. Subsequently, common variants that occurred in more than two sets were removed to obtain the final set of specific germline variants for each individual. The approach we described above allowed us to identify the cell clusters of each individual, as well as the sample-specific germline variants for the first time, thereby providing an efficient and robust method for downstream analysis such as doublet detection.

### Doublet detection

Definitions:

*K*: the number of clusters defined by the above steps.

*C*: the number of cells. Lower-case c is used for indexing and referring to a specific cell barcode.

*G*: the number of sample-specific germline variants sets. Lower-case *g* is used to represent all the specific germline variants in a cluster, equal to *G_i_* (*i* means cluster *i*). *g,c* means these cluster-specific germline variants with observed data in cell *c*.

*A*: allele counts. *A_g,c_* is a tuple of size 2 with the first number representing the number of reference alleles and the second number representing the number of alternate alleles observed in the cluster-specific germline variants in cell *c*.

We defined the initial clusters of cells or their corresponding cluster-specific germline variants. Our next objective was to identify doublets between each cluster.

For cluster-specific germline variants in each single cell, the observation of alternative allele counts follows a binomial distribution, with *p* being the probability of alternative allele frequency, *A_g,c,1_* being the alternative allele counts and *N* being the total counts for cluster-specific germline variants.

Equation (1):

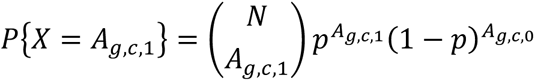

For a given cell, the probability of alternative alleles (*p*) or BAF at sample-specific germline variants can be modeled by a beta prior.

The Beta prior model for *p* can be tuned to reflect the relative prior plausibility of each *p*.

Equation (2): prior model:

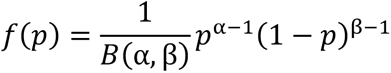

Upon observing data *X*=*x* where x∈ {0, 1, …, n}, the likelihood function of *p*, obtained by plugging *x* into the Binomial pmf Equation (1), we used Equation (3):

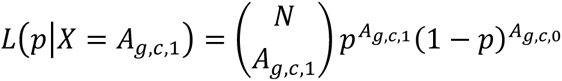

Via Bayes’ Rule, the conjugate Beta prior combined with the Binomial data model produce a Beta posterior model for posterior probability of cells belong to each cluster(*p*). Putting Equation (2) and Equation (3) together, the posterior pdf is descried by Equation (4):

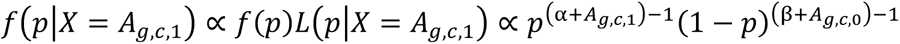

For each cluster, *α* and *β* can be estimated using the allele counts across all the sample-specific germline variants in one cluster.

*α* = 1 + mean of alternate counts across variants in *G_i_* of *K_i_*, and *β* = 1. (For cluster).

For a cell c belongs to first-fitted cluster *i* (*K_i_*), and *g*=*G*_i_, the *p_i_* was described by Equation (5):

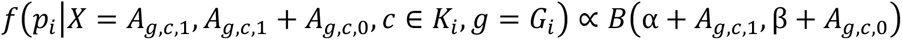

When the cell c belongs to first-fitted cluster i (K_i_) but g=G_j_, p_j_ was described by Equation (6):

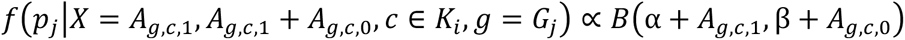

Then the *P_i_* was *p_1_* for the first-fitted cluster and the highest *P_j_* was selected as *p_2_* for the second fitted cluster. We estimate *p* as the mode of the posterior distribution. When the cell is a singlet, *p_1_* will be dramatically higher than any *P_j_*.

To give a reasonable threshold, we treated *p_1_*, *p_2_* as two-dimensional data. We treated *p_1_*>0.99 and *p_2_*<0.01 as “Singlet” and treated *p_1_*<0.8 or *p_2_*>0.2 as “Doublet” at the beginning, this cutoff can be set by users. To predict the class of unlabeled cells, we use the ‘KNeighborsClassifier’ method, which calculates the distance between the labeled cell point and all other unlabeled cell points in the dataset. The *k*=5 nearest neighbors are identified based on the distance metric, and the majority class of the *k*=5 nearest neighbors is used to predict the class of the unlabeled cells. It should be noted that if *p_1_* is greater than 0.99, we recommend not considering *p_2_*, as the value of *p*_2_ may be caused by potential somatic mutations.

### Demultiplexing pooled samples

After removing doublets that were identified in the previous step, individual cells are assigned to a cluster by the highest *p*. If cells are difficult to cluster initially under unsupervised clustering due to factors such as sequencing depth, highly imbalanced mixture proportions, and others, we will cluster the germline varinats and calculate probability values, refer to MitoSort ’--direct’ method. In cases where the exact number of pooled samples was unknown, we also provided the silhouette method to estimate the optimal number of clusters. The silhouette score reaches its global maximum at the optimal number of clusters, allowing for demultiplexing of pooled samples accurately.

### Simulation of *in silico* data

In this study, we simulated mixtures of mtscATAC-seq data obtained from the 10X Genomics platform using custom Python scripts. Following the alignment of reads to the human reference genome hg38 using 10X Genomics Cell Ranger ATAC(16), mitochondrial reads were merged into a bam file with cell barcodes prefixed to distinguish individuals. Next, reads were realigned using GATK(14). In total, we had 52283 cells across 8 samples. To assess the performance of MitoSort systematically, we generated synthetic mixtures into BAM files by random sampling using a range of parameter choices, including the number of reads mapped to mitochondrial genome (ranging from 250 to 2000), the number of pooled samples (ranging from 2 to 8), the number of cells for each sample (ranging from 100 to 600), and the percentage of doublets (1%, 5%, 10%, and 15%). Doublets were simulated by randomly choosing two cell barcodes and merging their read counts as a new cell. For each parameter set, simulations were repeated five times to account for variability.

### Benchmark against existing tools

We compared the performance of MitoSort against Souporcell (8), Vireo (7) and Freemuxlet (https://github.com/statgen/popscle) using simulated BAM file. Unless specified otherwise, default pipelines and recommended parameters were employed, and the number of threads utilized was set to 8. For Souporcell, souporcell_pipeline.py script was run within the singularity container with parameters “--no_umi True -- min_alt 2 --min_ref 2”. For Vireo, cellsnp-lite was used to genotype a list of variants in each cell with parameters “--minMAF 0.1 --minCOUNT 20 --UMItag None”. Subsequently, Vireo was utilized to demultiplex cells without any prior knowledge of genotypes. For Freemuxlet, the popscle dsc-pileup command was run within the Demuxafy (17)-provided singularity container to identify the number of reads from each allele at known variants with at leaset 1% minor allele frequency. Next, the popscle freemuxlet command was run to demultiplex cells. The effectiveness of each tool was assessed by comparing the classification outcomes with the true labels.

### Evaluation of MitoSort using cell hashing data

For multiplexed ASAP-seq dataset comprising NK cells from four donors (18), we obtained BAM file containing mitochondrial reads and HTO count data for ASAP5 library (GSM6413442) upon request from the corresponding author of the published paper. HTO count data containing hashtag reads were used to assign each cell a hashtag ID by the function HTODemux function within Seurat (19) package using default parameters. Subsequently, MitoSort was used to identify doublets and demultiplex samples based on the BAM file. Finally, a comparison was conducted between the donor assignment results obtained from MitoSort and those derived from the hashtag data.

For multiplexed DOGMA-seq dataset comprising activated and stimulated T cells from two healthy donors (20), we first downloaded its HTO count data (GSM6032896) and ATAC raw data (SRX14779147). HTO count data containing hashtag reads were used to assign each cell a hashtag ID by the function HTODemux function within Seurat (19) package. Raw reads in ATAC data were aligned to the human reference genome (hg38) using Cell Ranger ATAC (16). Based on the generated BAM file, we ran MitoSort pipeline, including MT reads realignment, variant calling, doublet identification and demultiplexing. Cell barcodes classified as singlets by hashtag and MitoSort were compared to evaluate the consistency of assignment results obtained from both approaches.

### Analysis of multiplexed Smart-seq3xpress data

We obtained raw data in the form of unmapped BAM files for two multiplexed Smart-seq3xpress datasets (21) from ERR8607752 and ERR8607757. These two datasets contain cells from two and three donors, respectively. For each dataset, the unmapped BAM file was further processed using zUMIs (https://github.com/sdparekh/zUMIs), which involved mapping to the human reference genome (hg38), and quantification of gene expression. Cells that passed the quality control criteria, as stated in the original paper, were retained for downstream analysis. To speed up the demultiplexing process, 20% of reads in BAM file were randomly sampled and used as input for MitoSort, Souporcell and Vireo, respectively. The donor assignment results from each tool were compared against the cell identities derived from dual indexes, serving as the ground truth labels, to assess the performance of each tool. Besides, we generated simulated BAM files with varying number of reads per cell and compared the runtime of each tool.

### Isolation of PBMCs and B cells

The study protocol was approved by the local ethics committee of Sun Yat-sen University (#2015/521), and written informed consent was obtained from all donors. All experiments and protocols for human studies involving human samples were performed in accordance with the relevant guidelines and regulations. To obtain peripheral blood mononuclear cells (PBMCs), the peripheral blood was diluted 1:1.5 in 0.9% physiological saline and cells were isolated with a Lymphoprep™ density gradient (density 1.077 g/mL) according to the manufacturer’s instructions (STEMCELL). B cells were isolated using Miltenyi CD19 MicroBeads (#130-050-301) as described by the manufacturer. Cells were counted using Trypan Blue exclusion under a hemocytometer.

### Single cell multi-omics library construction and sequencing

Single-cell multi-omics libraries were generated using the 10X Chromium Controller and the Chromium Next GEM Single Cell Multiome ATAC + Gene Expression kit according to the manufacturer’s instructions. DNA LoBind tubes (1.5 ml; Eppendorf) were used to wash cells in PBS and in downstream processing steps. To retain mtDNA, permeabilization was performed using LLL conditions, as described in mtscATAC-seq(3, 12) with 10 mM Tris-HCl pH 7.4, 10 mM NaCl, 3 mM MgCl_2_, 0.1% NP40 and 1% BSA. After washing, cells were fixed with 1% formaldehyde (ThermoFisher no. 28906) in PBS for 10 minutes at room temperature, followed by quenching with glycine solution to a final concentration of 0.125 M. After washing twice in PBS via centrifugation at 400g, 5 min, 4 °C, cells were treated with lysis buffer (10 mM Tris-HCl pH 7.4, 10 mM NaCl, 3 mM MgCl_2_, 0.1% NP40, 1% BSA) for 3 minutes, followed by adding 1 ml of chilled wash buffer (10 mM Tris-HCl pH 7.4, 10 mM NaCl, 3 mM MgCl2, 1% BSA) and inversion before centrifugation at 400g, 5 min, 4 °C. The supernatant was discarded, and cells were diluted in 1× Diluted Nuclei buffer (10X Genomics) before counting using Trypan Blue and a Countess II FL Automated Cell Counter. Briefly, after tagmentation, the cells were loaded on a Chromium Controller Single-Cell instrument to generate single-cell Gel Bead-In-Emulsions (GEMs) followed by PCR as described in the 10X protocol. The final libraries were quantified using a Qubit dsDNA HS Assay kit (Invitrogen) and a High Sensitivity DNA chip run on a Bioanalyzer 2100 system (Agilent). Libraries were sequenced on illumina NovaSeq 6000 at Berry Genomics.

### Analysis of multi-omics data of multiplexed B cells

Raw data of multiome were preprocessed and aligned to the human reference genome (hg38) using 10X Genomics Cell Ranger arc. The resulting BAM file from ATAC data was subjected to the MitoSort pipeline for doublet identification and sample demultiplexing. The cell-by-gene and cell-by-peak count matrices were further processed using the R package Seurat(19) and Signac (22). High-quality cells with less than 5000 RNA molecules, mitochondrial RNA content <10%, 200-3500 expressed genes, 2000-50000 fragments, TSS enrichment score >= 5, nucleosome signal <1, and the percentage of reads in peaks >= 50 were retained for downstream analysis. Subsequently, weighted nearest neighbor (WNN) analysis was performed on both RNA and ATAC-seq modalities, resulting in 10 clusters. Cells were then annotated into four cell types using the following marker genes: B cells (BANK1, MS4A1); Plasma (MZB1); T cells (CD3D, CD3E); monocytes (CD14, S100A9). After removing cross-genotype doublets assigned by MitoSort, we obtained a total of 12943 B cells. Next, WNN analysis was applied to the subclustering analysis of multiome data of B cells, which yielded 5 clusters. B cells were further divided into two subtypes using the following marker genes: naive B cells (TCL1A, IL4R); memory B cells (CD27). Based on the donor origins of cells obtained from MitoSort, the percentage of each cluster in each sample was calculated.

## RESULTS

### Overview of MitoSort

The MitoSort pipeline is mainly composed of four steps: MT reads realignment, variant calling, doublet identification and demultiplexing (Figure 1b). Briefly, to reduce false-positive variants resulting from STAR (23) alignments artifacts, MT reads are realigned with GATK(14) to facilitate accurate variant calling. VarScan (15) is then used to identify putative single-nucleotide polymorphisms (SNPs) in mtDNA. An SNP-by-cell matrix is generated after filtering low-frequency variants. By the initial K-means clustering, sample-specific germline variants are identified, allowing for the precise clustering of cells based on unique mitochondrial haplotypes. To detect cross-genotype doublets, we model a cell’s allele counts as being drawn from a beta-binomial distribution, with parameters derived from one or two clusters. After the removal of doublets, singlet cells are assigned to their respective donor of origin by assessing the probabilities associated with each cluster. Additionally, our model estimates the most likely number of pooled individuals, which is helpful when clinical information is lost (Methods). It only takes 10 minutes for demultiplexing a mixture of ∼10000 cells by MitoSort.

### Evaluation of MitoSort using *in silico* data

To assess the performance of MitoSort, we first used in silico data with known cell origins to evaluate its accuracy and efficacy. In order to generate synthetic datasets, we collected eight published mtscATAC-seq datasets from genetically distinct donors (detailed information is shown in Supplementary Data Table1). Current computational tools are mainly designed for demultiplexing scRNA-seq data using nuclear genome variants. Although some of them can be adapted to process mtscATAC-seq data, their performance on mtscATAC-seq data has not been systematically evaluated thus far. We applied MitoSort along with three existing computational tools, including Souporcell (8), Vireo (7) and Freemuxlet (5), to the synthetic datasets and compared their performance. As expected, the results demonstrated that both Souporcell and Vireo require higher sequencing depth to assign cells. Due to the inherent sparsity of genome coverage in mtscATAC-seq data, a significant proportion of cells cannot be assigned to their respective donors (Figure 2a). Although the proportion of unassigned cells decreased when the number of mapped reads per cell ranged from 10000 to 50000 (8-mixed samples, 100 cells per sample and 8% of cells as doublets), it still remained higher than that achieved by MitoSort. Besides, despite consuming more than eight times the running time of MitoSort (Figure 2b), these tools fail to accurately identify cross-genotype doublets and assign singlets to their corresponding original donors (Figure 2c-d, Supplementary Data Figure 1). These results revealed the inadequate performance of previous computational tools when applied to mtscATAC-seq data. MitoSort fills this critical void by offering improved functionality in demultiplexing mtscATAC-seq data.

**Figure 2.**
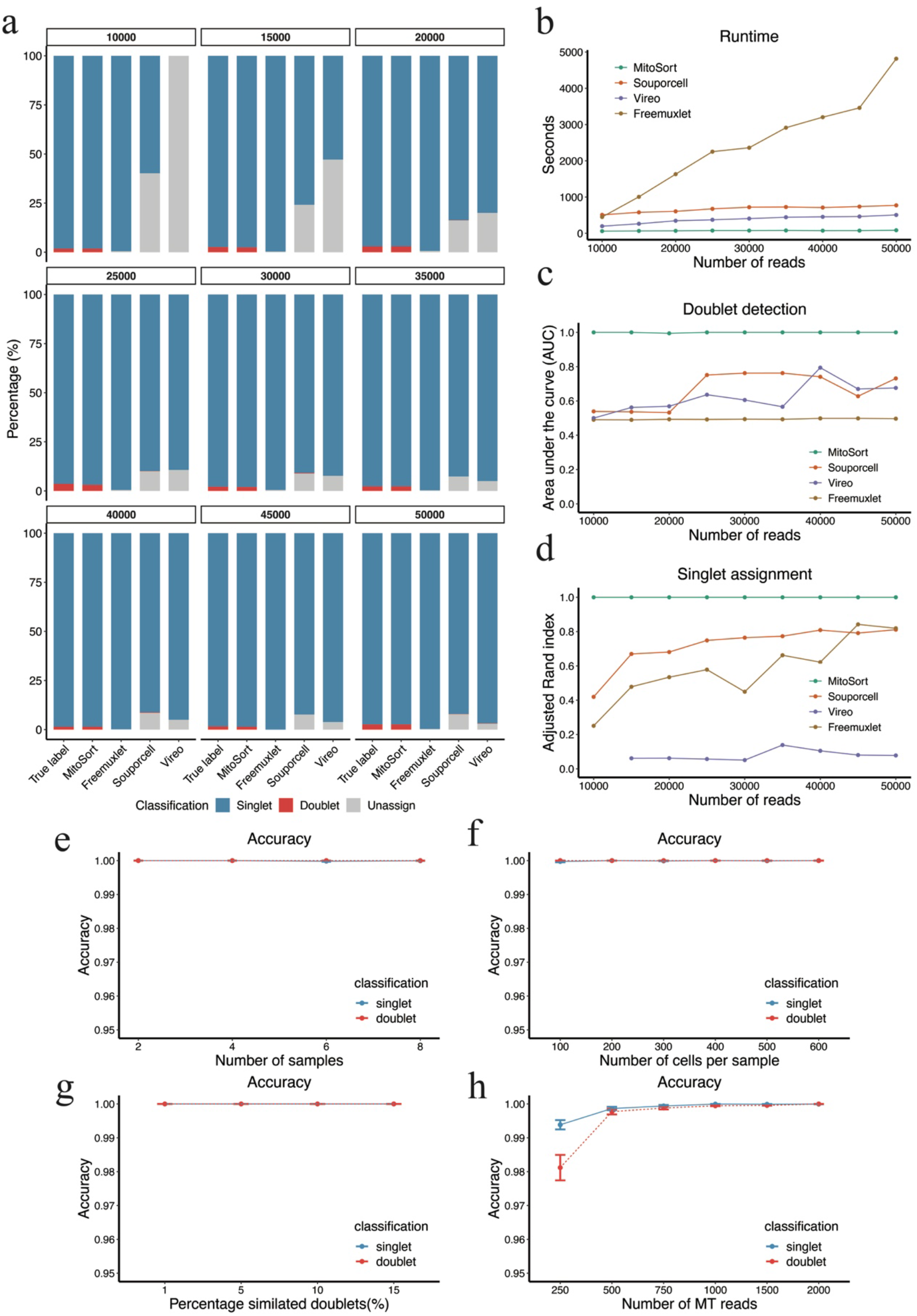
Evaluation of MitoSort using *in silico* data. **a**, Percentage of cell assignment of each tool on simulated data containing eight pooled samples, each with 100 cells and varying numbers of reads per cell. **b,** Runtime of each tool on simulated data containing eight pooled samples, each with 100 cells and varying numbers of reads per cell. **c,** The area under the curve (AUC) for doublet identification of each tool on simulated data containing eight pooled samples, each with 100 cells and varying all reads per cell. **d,** The Adjusted Rand index (ARI) between the true and inferred singlet assignment for each tool on simulated data containing eight pooled samples, each with 100 cells and varying all reads per cell. **e-h,** Systematic assessment of MitoSort performance on simulated data using a range of parameter choices, five replicate runs are used to ensure the consistency of the results. **e,** Classification accuracy of MitoSort on simulated datasets comprising 2-8 mixed samples, each with 500 cells, an 8% doublet rate, and 2000 MT reads per cell. **f,** Classification accuracy of MitoSort on simulated datasets containing eight pooled samples, each with varying numbers of cells, an 8% doublet rate and 2000 MT reads per cell. **g,** Classification accuracy of MitoSort on simulated datasets containing eight pooled samples, each with 500 cells, 2000 MT reads per cell and varying doublet rates. **h,** Classification accuracy of MitoSort on simulated datasets containing eight pooled samples, each with 500 cells, an 8% doublet rate, and varying numbers of mitochondrial (MT) reads per cell.

We next further assessed the impact of factors affecting the classification performance of MitoSort. Among these factors, the number of pooled samples, the number of cells for each sample, and the percentage of doublets had little effect on the classification performance of MitoSort (Figure 2e-g, Supplementary Data Figure 2a-f). We consistently obtained high accuracy of origin assignment and doublet identification for 2- to 8-mixed samples (500 cells per sample, 2000 mtDNA reads per cell, and 8% of cells as doublets) (Figure 2e). Besides, when we synthetically mixed 8 samples (2000 mtDNA reads per cell and 8% of cells as doublets) and varied the number of cells from 100 to 600 per sample, we found the true positive rate (TPR) of both donor assignment and doublet identification was consistently achieved at a rate of 1, even when the number of cells was as low as 100 (Supplementary Data Figure 2c). Only a small number of singlets were erroneously classified as doublets, constituting less than 0.3% of all cases (Supplementary Data Figure 2d). In addition, MitoSort achieved perfect performance regardless of the percentage of simulated doublets (Figure 2g, Supplementary Data Figure 2e-f). We found the performance of MitoSort was only sensitive to the sequencing depth of mitochondrial genome (Figure 2h). However, even with as few as 250 mapped reads on the mitochondrial genome, MitoSort achieved an accuracy exceeding 97%. Specifically, an increase in the number of mapped reads (ranging from 250 to 2000) resulted in a higher true positive rate (TPR) and a lower false discovery rate (FDR), leading to improved accuracy (Figure 2h, Supplementary Data Figure 2g-h). It is worth noting that when the number of reads mapped to the mitochondrial genome reached approximately 1000 (representing roughly 4X), MitoSort was able to achieve perfect cell assignment (100% accuracy) to their respective donors and accurately detect cross-genotype doublets. Collectively, these data show that MitoSort is a highly accurate and scalable method for demultiplexing scATAC-seq data derived from distinct individuals and identifying cross-genotype doublets using mtDNA germline variants.

### Evaluation of MitoSort using cell hashing data

Cell hashing with oligo-tagged antibodies allows for the assignment of sample origin to cells in mtscATAC-seq datasets based on the counts of hashtag oligonucleotides (HTOs) (24). Using the HTO-based assignment as ground truth, we next evaluated our method with two published experimentally multiplexed datasets using cell-hashing method. For ASAP-seq dataset comprising NK cells from four donors(18), MitoSort demonstrated high consistency (∼98%) in donor identity assignment (Figure 3a). Singlets originating from the same donor exhibited identical MT genotypes (Figure 3b-c). Compared to hashtag counts, the bimodal distributions of allele frequency for donor-specific mitochondrial germline variants exhibited more pronounced disparities among different singlet groups (Figure 3d-e). demonstrating that mtDNA variants can serve as a more computationally efficient and robust feature for donor assignment. For doublet identification, we observed that hashtag-based approach identified more doublets than MitoSort, resulting in a notable proportion of cells with inconsistent singlet/doublet classifications between the two strategies (Figure 3a). Upon examining of the mitochondrial (MT) haplotypes of these discordant cells, we observed that cells that were classified as doublets solely by MitoSort indeed contained multiple MT genotypes, whereas cells that were classified as doublets solely by hashtag-based approach contain only one MT genotype (Figure 3f). We next compared the numbers of chromatin fragments in the three doublet groups with that of the singlet group. We found the doublets defined by the two methods exhibited a significantly higher sequencing depth compared to the singlet group, while the doublets defined only by MitoSort (Doublet-Singlet group) showed a slightly higher sequencing depth (Figure 3g). Encouraged by the clarity of MT genotype (Figure 3f), we further examined the original counts of hashtags for the cells in the Doublet-Singlet group. We found the HTO demultiplexing tool tended to assign cells with higher HTO counts as doublets (Supplementary Data Figure 3-2 a-b). The cells in the Doublet-Singlet group have two HTO tag called, but with lower counts, leading to the false negative identification of doublets (Supplementary Data Figure 3-2 c-e, 3-3 a-c). Interestingly, the number of chromatin fragments in doublets called only by hashtag-based approach (Singlet-Doublet group) is also higher than that of the singlet group (Figure 3g), suggesting that these cells may be doublets from the same genotype, which cannot be distinguished by MitoSort. However, the proportion of doublet contributing by the same genotypes is estimated to be 1.8 % of total cells (250 cells out of a total of 13919) according to the number of pooled donors. The proportion of observed cells in Singlet-Doublet group was about 8% (1108 cells), which is much higher than the expected number, indicating there are false positive calling by hashtag-based approach (Supplementary Data Figure 3-2 c-e, 3-3 a-c). In addition to its accuracy, MitoSort also showed higher sensitivity compared to the hashtag-based approach. MitoSort was able to assign donor and singlet/doublet status for 99.1% of the cells, compared to only 62.4% achieved by hashtag (Supplementary Data Figure 3-1 a-b). These findings indicated that the performance of demultiplexing by exogenous hashtags were heavily dependent on the outcome of hash-tagging experiment, while endogenous germline mitochondrial variants were more stable and could be used for evaluating of experimental efficiency and quality control measures.

**Figure 3.**
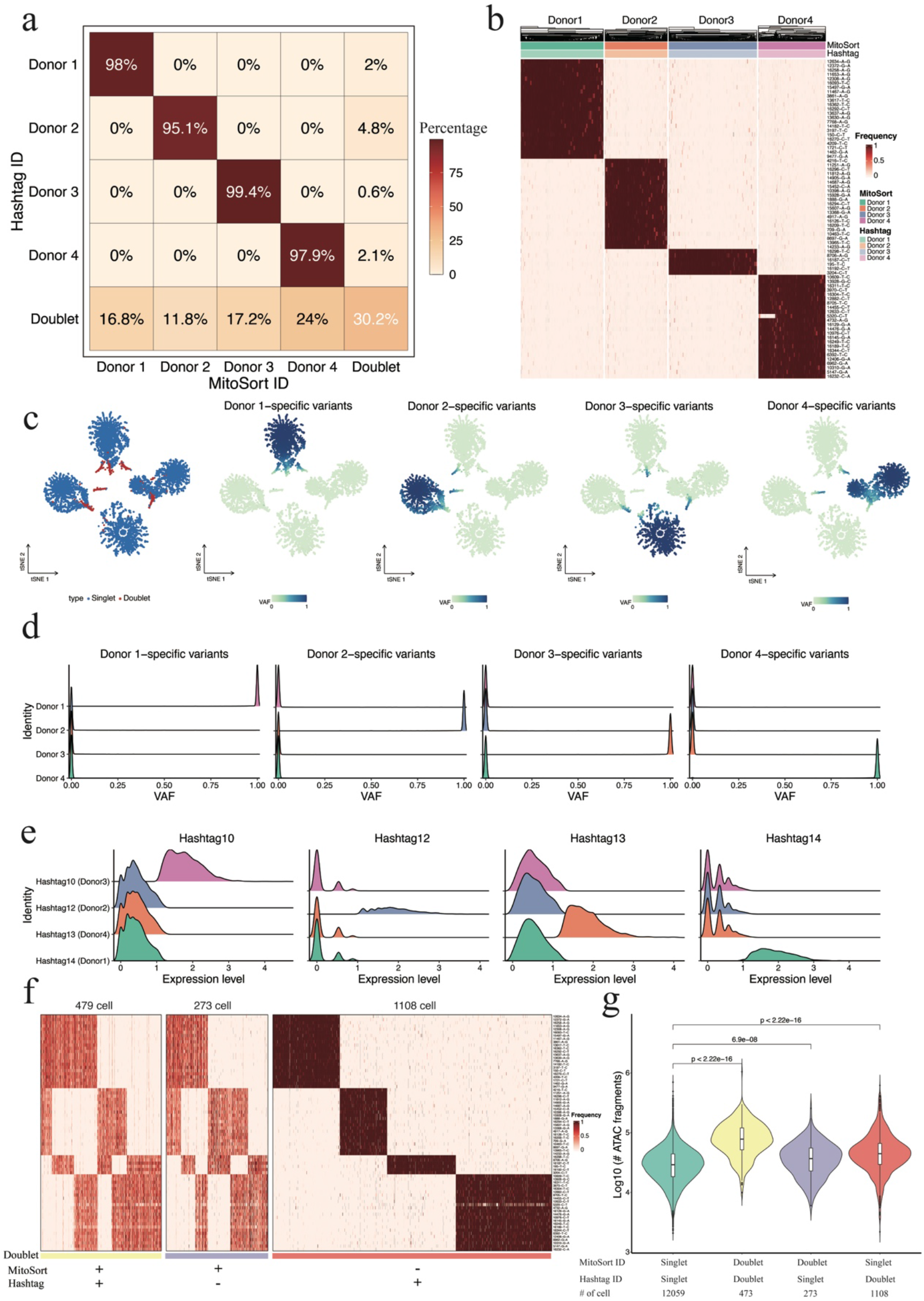
Evaluation of MitoSort using cell hashing data. **a**, Percentage of barcodes shared between hashtag-based (rows) and MitoSort-based (columns) assignments in multiplexed ASAP-seq data consisting of four donors. **b,** Heatmap showing the allele frequency of sample-specific mitochondrial germline variants (rows) across singlets (columns) in multiplexed ASAP-seq data. Top legend shows the donor assignments of MitoSort and hashtag-based approach. **c**, t-SNE plots of cell barcodes by frequency of sample-specific mitochondrial germline variants, colored by singlet/doublet classification (left) or by average frequency for the donor-specific mitochondrial germline variants (four right panels). **d,** Distributions of frequency of sample-specific germline variants for each donor across four singlet groups assigned by MitoSort. **e,** Distributions of counts for each hashtag across the four hash IDs. **f,** Heatmap showing the allele frequency of sample-specific mitochondrial germline variants (rows) across cells (columns) for concordant and discordant doublets. **g,** Distribution of the number of ATAC fragments per cell in four groups with concordant or discordant assignments (the Wilcoxon Rank Sum Test).

MitoSort also achieved great performance on the DOGMA-seq dataset which consisted of activated and stimulated T cells from two healthy donors (20). Notably, when the sequencing depth of mitochondrial genome was not less than 4X, MitoSort consistently achieved a high degree of accuracy in assigning cellular origins, with an accuracy rate of no less than 98% (Supplementary Data Figure 3-3 d-e). Collectively, these results further suggest the robustness of MitoSort in assigning cells from different donors and identifying cross-genotype doublets using mtDNA germline variants.

### Application of MitoSort to demultiplex pooled full-length scRNA-seq

MitoSort is designed and validated for mtscATAC-seq data which captures the complete mitochondrial sequences at high coverage. Previous studies have demonstrated that mitochondrial variants can be also captured in single-cell RNA sequencing protocols with high coverage (25, 26). We next evaluated the coverage and capture rate of mitochondrial genome in eight single-cell sequencing techniques (Figure 4a-b). Compared to droplet-based single-cell RNA sequencing techniques that typically sequence the 3’ or 5’ ends of RNA molecules, full-length single-cell RNA-seq protocols offer improved coverage of mitochondrial transcripts and facilitate variant calling for larger parts of the mtDNA genome (Figure 4a). In each full-length scRNA-seq dataset, approximately over 90% of cells achieved an average sequencing depth of more than 5X across the entire mitochondrial genome (Figure 4b, Supplementary Data Figure 4a). This demonstrates the feasibility of utilizing MitoSort to demultiplex full-length scRNA-seq data as well.

**Figure 4.**
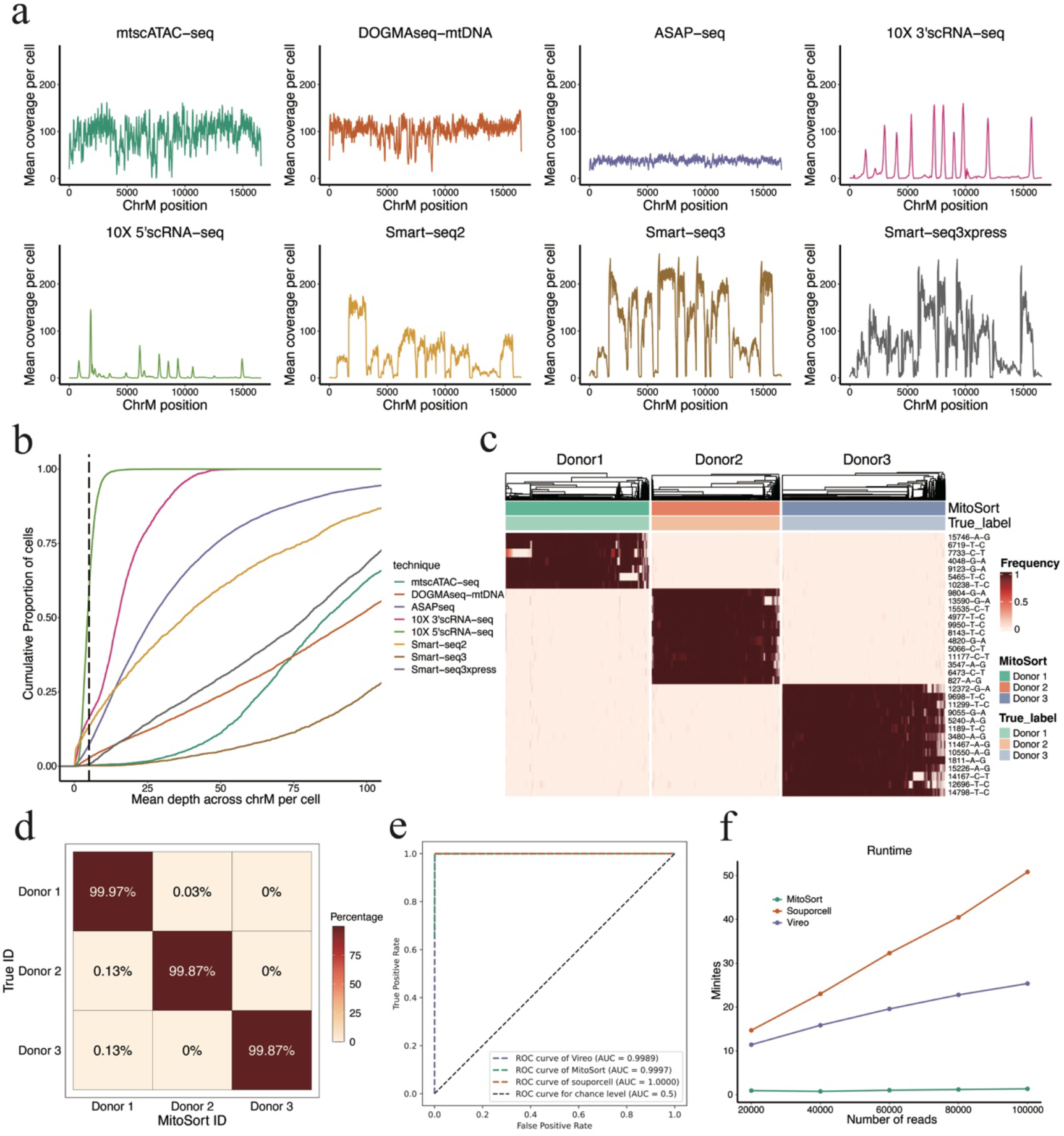
Extension of MitoSort to full-length scRNA-seq data. **a**, Coverage comparison across three scATAC-seq datasets and five scRNA-seq datasets. Droplet-based scRNA-seq shows uneven distribution of reads across the mitochondrial genome while others are generally more well-covered. **b,** An Empirical Cumulative Distribution Function (ECDF) plot showing average sequencing depth across mitochondrial genome per cell for eight datasets using different sequencing techniques. **c,** Heatmap showing the allele frequency of sample-specific mitochondrial germline variants (rows) across MitsoSort-assigned singlets (columns) in multiplexed Smart-seq3xpress dataset. Top legend shows the donor assignments obtained from MitoSort and dual indexes. **d,** Percentage of barcodes shared between dual indexes-based (rows) and MitoSort-based (columns) assignments in multiplexed Smart-seq3xpress data consisting of three donors. **e,** Receiver operating characteristic (ROC) curves for singlet assignment of each tool on multiplexed Smart-seq3xpress data consisting of three donors. **f,** Runtime of each tool on simulated Smart-seq3xpress data containing three donors, each with 200 cells and varying numbers of reads per cell.

We next evaluated the performance of MitoSort using the Smart-seq3xpress dataset, which consisted of cells from three donors (21). The classification results obtained by MitoSort were highly consistent with true donor identities derived from dual indexes (Figure 4d), with cells assigned to the same donors sharing identical mitochondrial haplotype (Figure 4c). We also benchmarked our method against Souporcell and Vireo, two of the best computational tools for demultiplexing pooled single-cell RNA-seq data using genome variants. MitoSort demonstrated comparable performance of accuracy in singlet assignment to the tools specifically developed for scRNA-seq data (Figure 4e) and showed higher accuracy in doublet identification (Supplementary Data Figure 4b). Furthermore, MitoSort exhibited better computational efficiency compared to other tools, with superior processing speed when applied to the same dataset (Figure 4f). We obtained similar results when applying MitoSort to another Smart-seq3xpress dataset comprising cells from two donors (Supplementary Data Figure 4c-d). Collectively, these results demonstrate the versatility of MitoSort. It can be employed for accurately demultiplexing pooled datasets derived from variant single-cell genomics libraries, provided that the mitochondrial sequence can be appropriately readout.

### Application of MitoSort in a multi-omics study voiding batch effects

Due to the high cost of single cell genomics approaches, conducting technical replicates is often unfeasible for most of the current studies. However, batch effects can diminish the sensitivity of identifying true biological signals and may introduce artifacts. Demultiplexing samples within the same experimental run can effectively reduce artifacts arising from sample preparation and library construction, thereby enhancing the accuracy and reliability of downstream analyses.

We conducted a case study in which we multiplexed B cells from eight individuals to profile gene expression, chromatin accessibility, and mitochondrial DNA (Figure 5a). The scATAC-seq library contained an average of 25% mtDNA reads per cell, which allowed for accurate mtDNA variant calling and demultiplexing (Supplementary Data Figure 5a). Using MitoSort, we could clearly classify a total of 16472 cells into eight singlet populations, as well as doublet groups based on the allele frequency of unique mtDNA germline variants for each individual (Figure 5b). 6.4% of cell barcodes were identified as cross-genotype doublets involved more than one MT genotype (Figure 5b, Supplementary Data Figure 5b-c). Further analysis showed an expected higher number of RNA molecules and chromatin fragments in these doublets (Supplementary Data Figure 5d). Following the removal of low-quality cells and cross-genotype doublets, we obtained 12943 high-quality B cell singlets. High-dimensional clustering of multi-omic profiles identified 5 clusters, which could be annotated as naïve B cells and memory B cells based on known markers (Figure 5c, Supplementary Data Figure 5e-f). Among the 8 individuals, we found the proportions of each B cell subtype are significantly variant (Figure 5d-e). Using mtDNA SNP information, we can match donors with MitoSort-assigned labels. According to the clinical records, the individuals with lower proportion of memory B cells had received rituximab (RTX) treatments, which was consistent with the function of RTX (27). It suggested that the observed differences in cell composition reflects true biological differences. Collectively, these data demonstrated that MitoSort facilitated accurately demultiplexing samples of individuals under different pathologies, thereby effectively distinguishing biological difference from technical batch effects.

**Figure 5.**
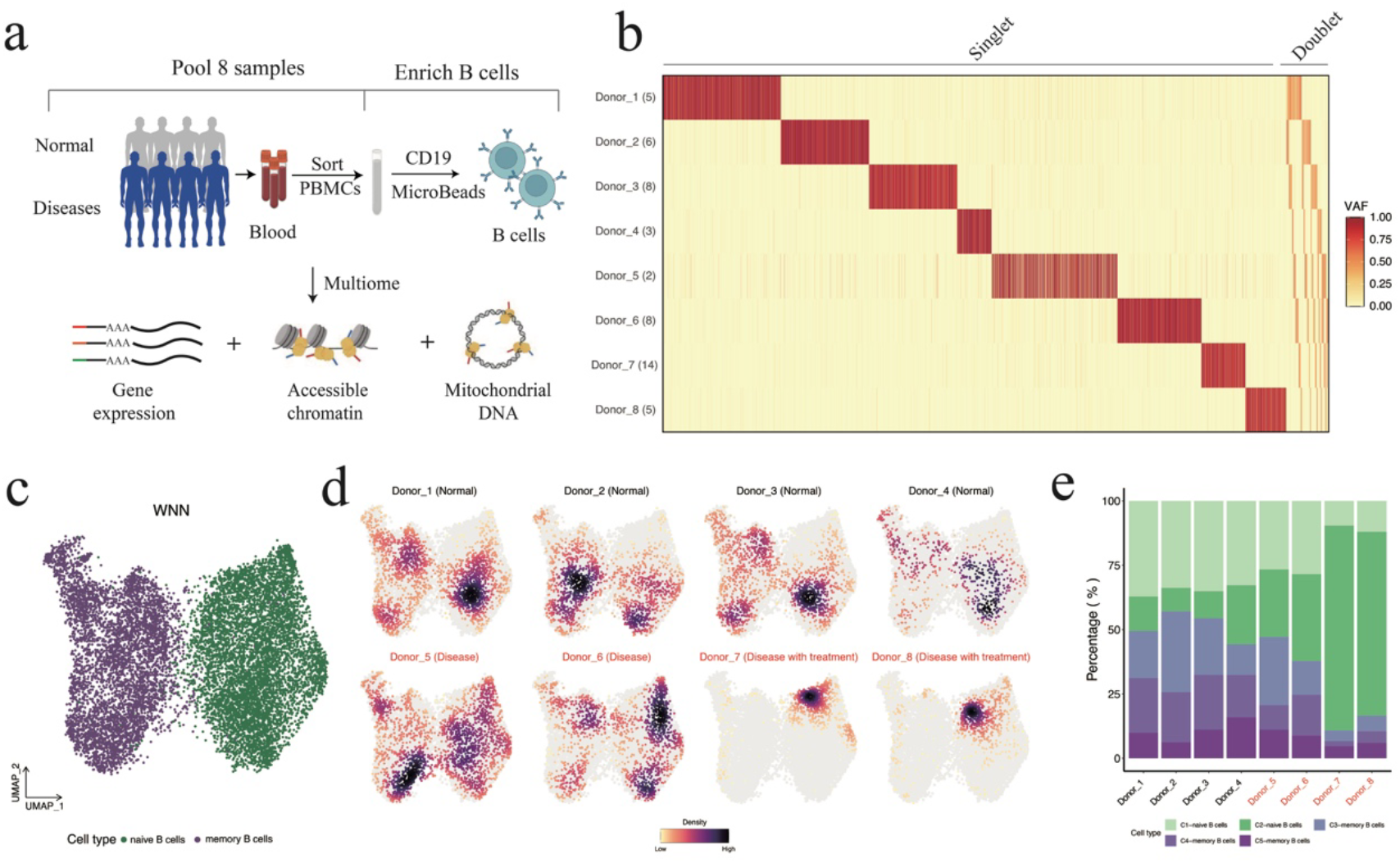
Multi-omics analysis based on MitoSort identified depletion of B cells RTX-treated patients. **a**, Schematic of experimental design. Peripheral blood mononuclear cells (PBMCs) from four normal individuals and four patients with antoimmune diseases were collected and pooled. Next, B cells were enriched to profile gene expression, chromatin accessibility, and mitochondrial DNA. **b,** Heatmap showing the allele frequency of aggregated sample-specific mitochondrial germline mutations (rows) across cells (columns) sorted by MitoSort assignment. The number in the parentheses of each row name represents the number of sample-specific mitochondrial germline mutations for this sample. **c,** UMAP of weighted-nearest neighbor (WNN) graph for RNA and ATAC-seq modalities showing cell type annotation for each cell. **d,** UMAP of weighted-nearest neighbor (WNN) graph for RNA and ATAC-seq modalities showing cells per individual colored by cell number densities. Disease status and clinical information for each individual are shown in subtitles. **e,** Bar plot showing the percentage of each cell cluster in each individual.

## DISCUSSION

In conclusion, the unique properties of mitochondrial DNA (mtDNA) make it a more efficient and robust choice for endogenous genetic barcodes in sample demultiplexing and the detection of cross-genotype doublets. However, only a few software options are currently available for this purpose. MitoSort fills this gap by providing an accurate and scalable method for demultiplexing pooled single-cell genomics data (mtscATAC-seq, ASAP-seq, DOGMA-seq, Smart-seq) across donors and identifying cross-genotype doublets. Nonetheless, the matrilineal inheritance of mitochondria poses a limitation to this approach, as it is challenging to distinguish individuals with haploidentical mtDNA.

Our systematical comparison revealed that all the currently available methods for doublet detection have their own limitations and bias. Profiling-based methods, which rely on the transcriptional profiles for doublet detection (28), may not be effective in cases where the pooled cells lack cell type diversity. Genotyping-based methods, including MitoSort, fail to detect doublets when the cells originate from the same individual. Experimental-based methods for doublet detection rely on the labeling efficiency. However, the advantage of genotyping-based methods is that as the number of pooled samples increases, the probability of doublet from the same individual decreases. We prospect that genotyping-defined doublets could serve as valuable positive controls to guide the refinement of doublet detection through profiling-based methods, ultimately enhancing the accuracy of doublet identification in future studies.

## DATA AVAILABILITY

Single-cell multiome data generated in this study has been deposited to National Genomics Data Center (NGDC) under the accession number PRJCA016782. Access to the data can be obtained upon request according to the police on human resources. MitoSort pipeline and scripts for reproducing the study are available at GitHub (https://github.com/tangzhj/MitoSort).

## AUTHOR CONTRIBUTIONS

J. X. designed and conceived the study. Z.T., P.S. collected and analyzed the data. X.L., S.L. and Y.X. supported the data analysis. Y.S. and W.Z. collected samples. W.Z. generated the single cell multi-omic library. J. X., Z.T. and P.S. wrote the manuscript with inputs from all authors. All authors read and approved the final manuscript.

## ACKNOWLEDGEMENTS

We thank Timo Rückert for sharing raw data of ASAP5 library with us.

---

**Supplementary Data Figure 1.**
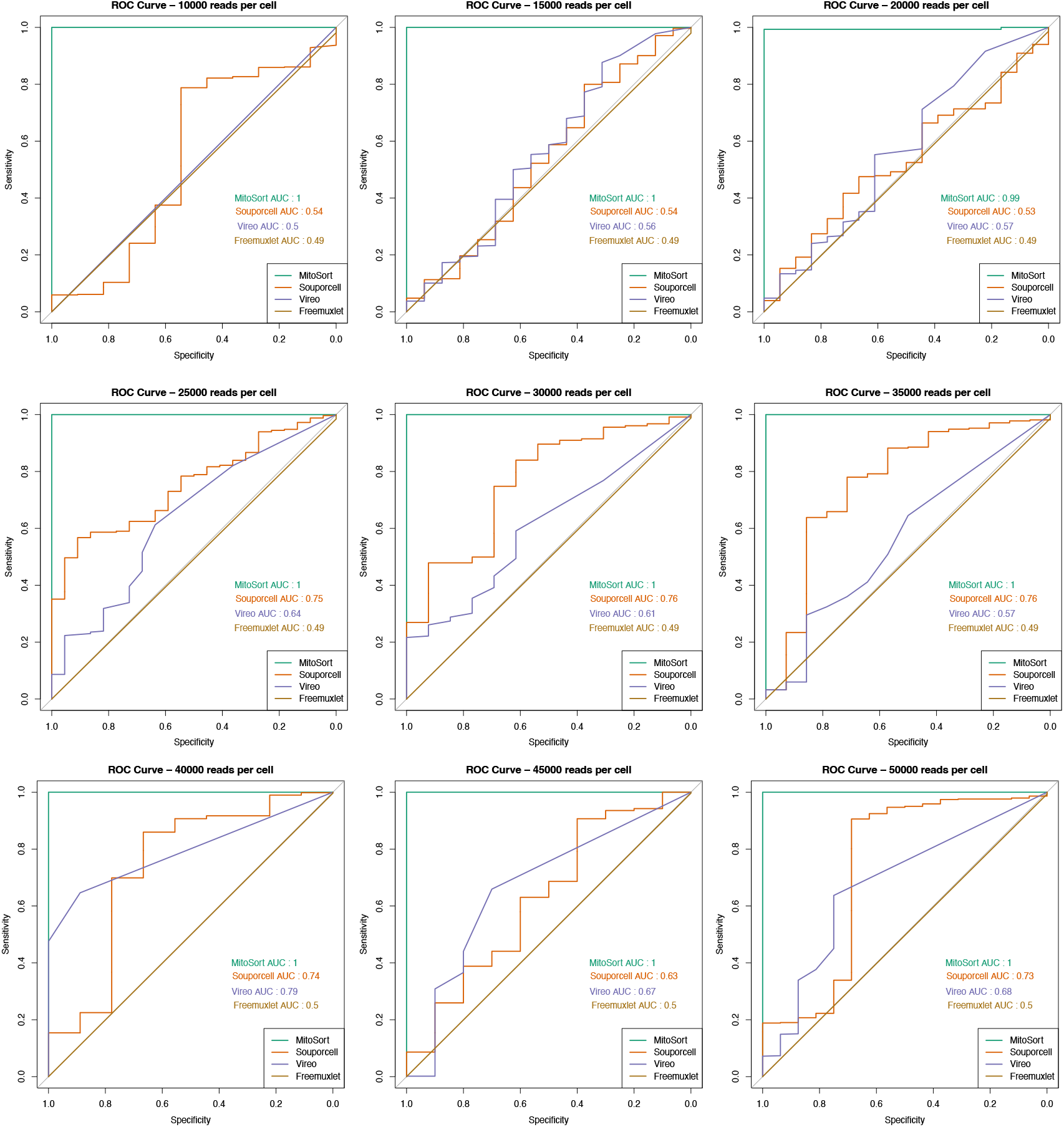
Receiver operating characteristic (ROC) curves for doublet identification of each tool on simulated data containing eight pooled samples, each with 100 cells and varying all reads per cell.

**Supplementary Data Figure 2.**
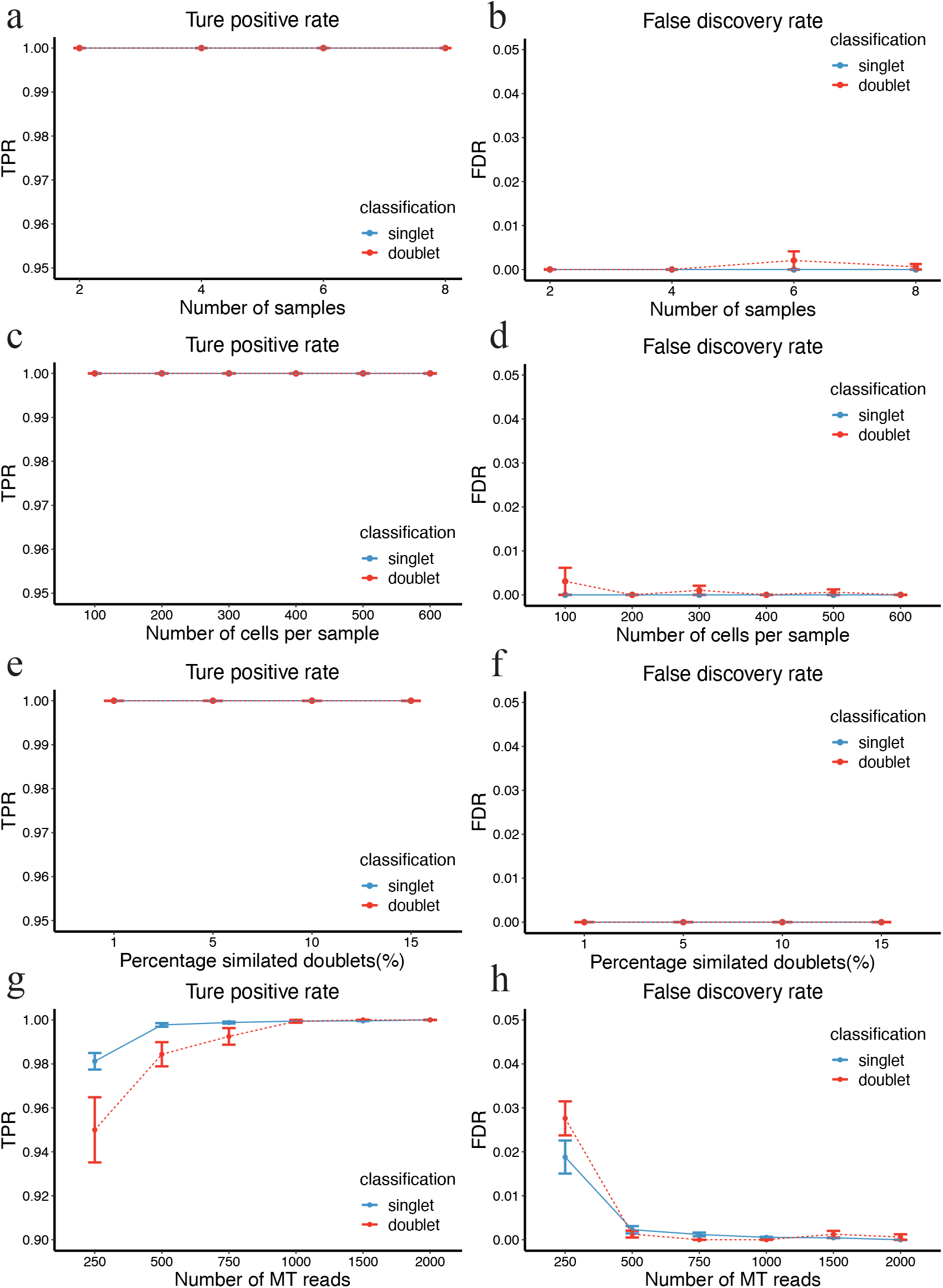
Systematic assessment of MitoSort performance on simulated data using a range of parameter choices. **a-b**, The true positive rate (TPR) and false discovery rate (FDR) of donor assignment and doublet identification by MitoSort on simulated datasets comprising 2-8 mixed samples, each with 500 cells, an 8% doublet rate, and 2000 MT reads per cell. **c-d,** The true positive rate (TPR) and false discovery rate (FDR) of donor assignment and doublet identification by MitoSort on simulated datasets containing eight pooled samples, each with varying numbers of cells, an 8% doublet rate and 2000 MT reads per cell. **e-f,** The true positive rate (TPR) and false discovery rate (FDR) of donor assignment and doublet identification by MitoSort on simulated datasets containing eight pooled samples, each with 500 cells, 2000 MT reads per cell and varying doublet rates. **g-h,** The true positive rate (TPR) and false discovery rate (FDR) of donor assignment and doublet identification by MitoSort on simulated datasets containing eight pooled samples, each with 500 cells, an 8% doublet rate, and varying numbers of mitochondrial (MT) reads per cell.

**Supplementary Data Figure 3-1.**
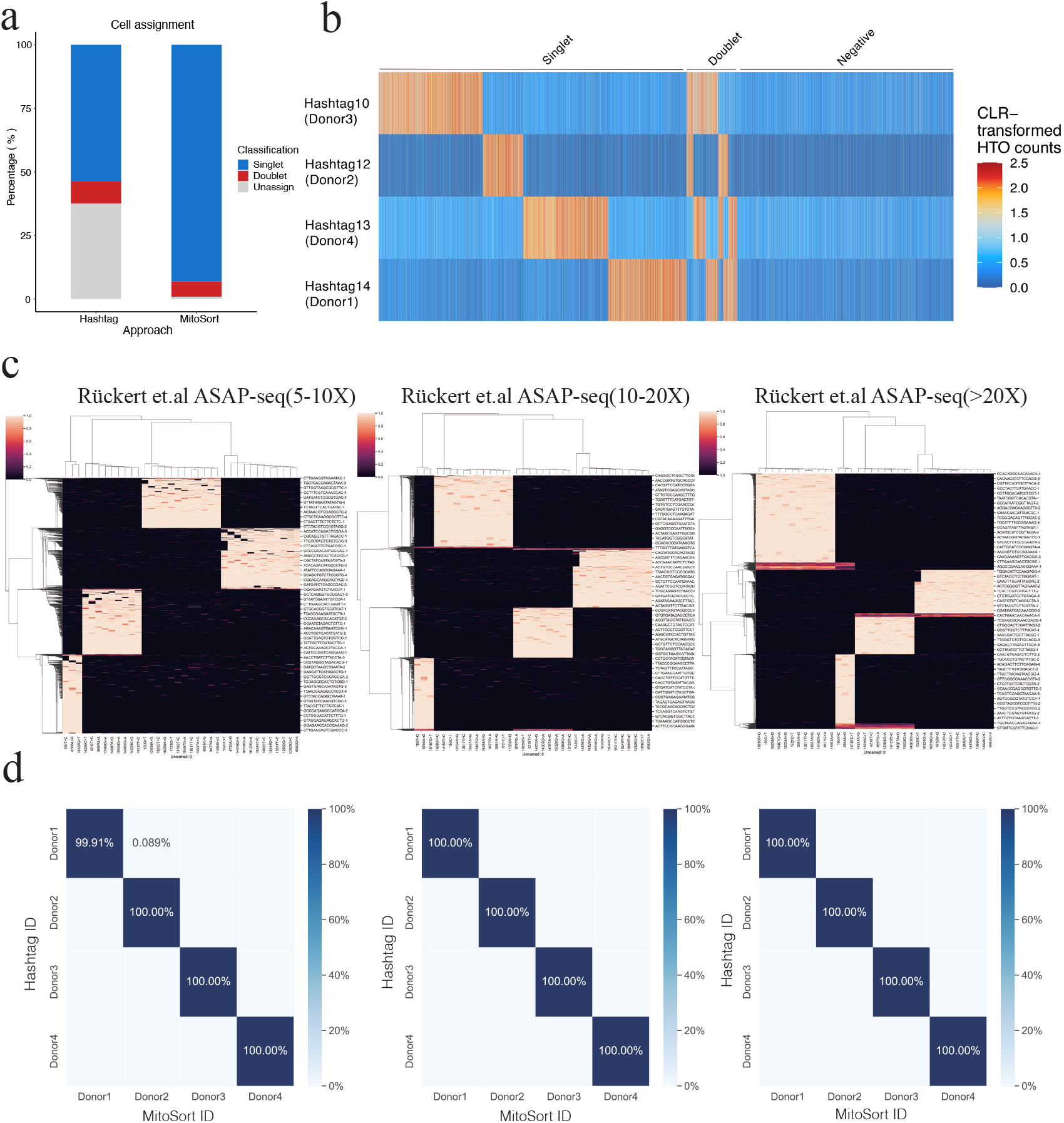
Evaluation of MitoSort using cell hashing data. **a**, Percentage of cell assignment of MitoSort and hashtag-based approach on multiplexed ASAP-seq data consisting of four donors. **b,** Read counts for each hashtag (rows) across cells (columns) sorted by their HTODemux classification. **c,** Heatmap showing the allele frequency of sample-specific mitochondrial germline mutations (columns) across cells (rows) with different sequencing depth of mitochondrial genome (5-10X, 10-20X and >20X) in multiplexed ASAP-seq data. **d,** Percentage of barcodes with different sequencing depth of mitochondrial genome (5-10X, 10-20X and >20X) shared between hashtag-based (rows) and MitoSort-based (columns) assignments in multiplexed ASAP-seq data.

**Supplementary Data Figure 3-2.**
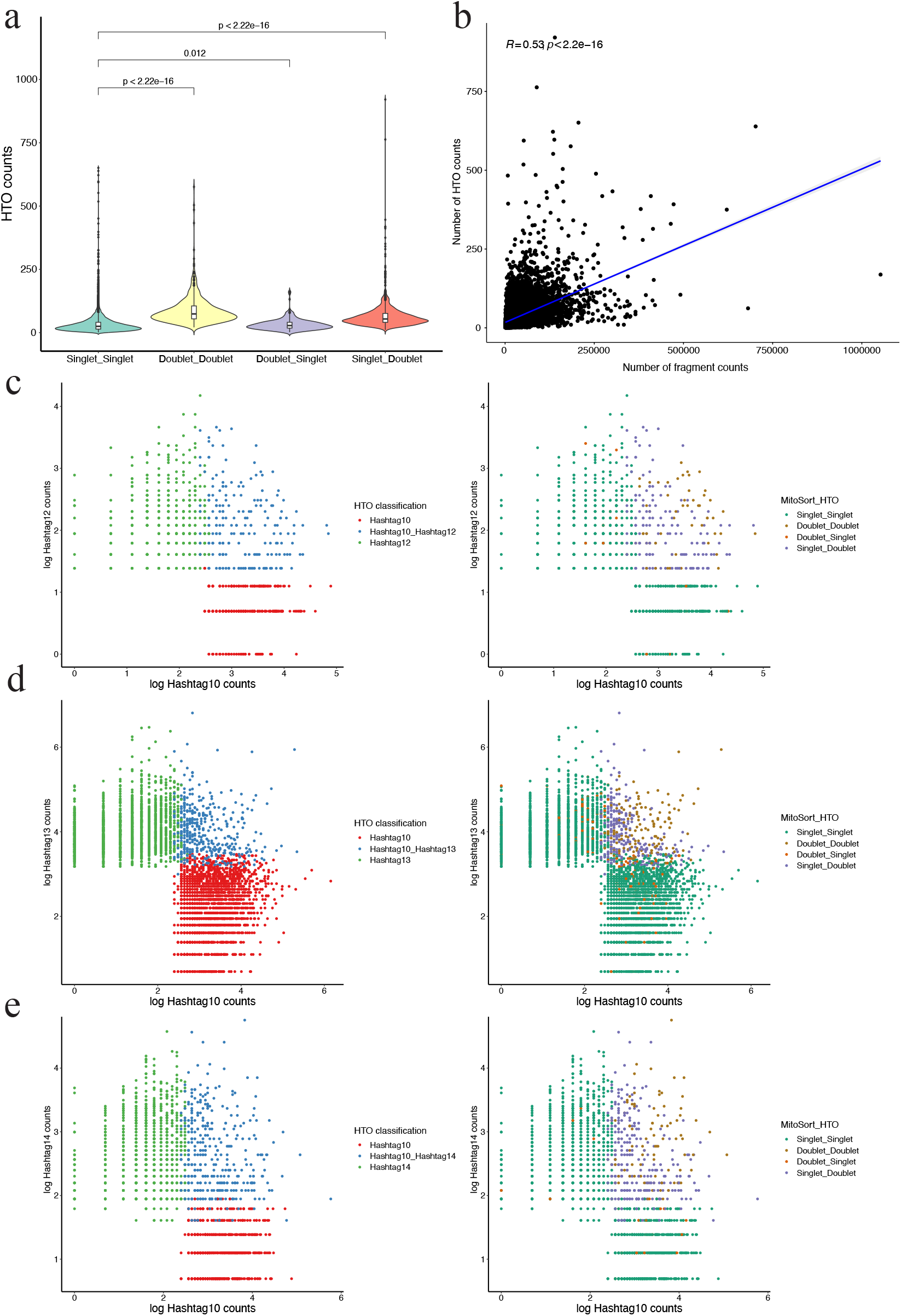
Evaluation of MitoSort using cell hashing data. **a**, Distribution of hashtag counts in four groups with concordant or discordant assignments (the Wilcoxon Rank Sum Test). **b,** Correlation of the number of ATAC fragments and hashtag counts. **c-e,** Scatter plots showing raw counts for pairs of hashtags colored by HTO-based assignment (left) or by combined assignment from MitoSort and HTO-based approach (right).

**Supplementary Data Figure 3-3.**
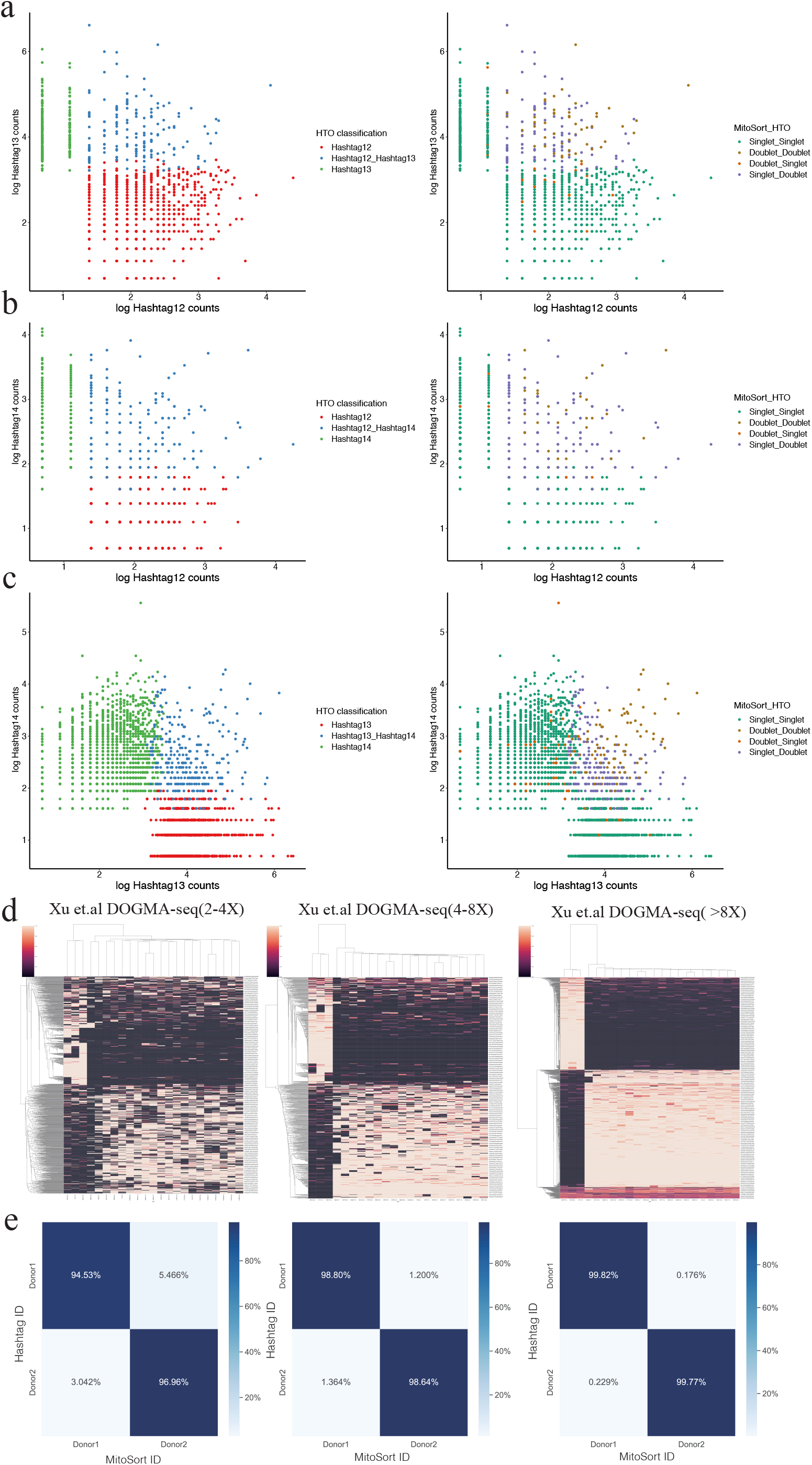
Evaluation of MitoSort using cell hashing data. **a-c**, Scatter plots showing raw counts for pairs of hashtags colored by HTO-based assignment (left) or by combined assignment from MitoSort and HTO-based approach (right). d, Heatmap showing the allele frequency of sample-specific mitochondrial germline mutations (columns) across cells (rows) with different sequencing depth of mitochondrial genome (2-4X, 4-8X and >8X) in multiplexed DOGMA-seq data. **e,** Percentage of barcodes with different sequencing depth of mitochondrial genome (2-4X, 4-8X and >8X) shared between hashtag-based (rows) and MitoSort-based (columns) assignments in multiplexed DOGMA-seq data.

**Supplementary Data Figure 4.**
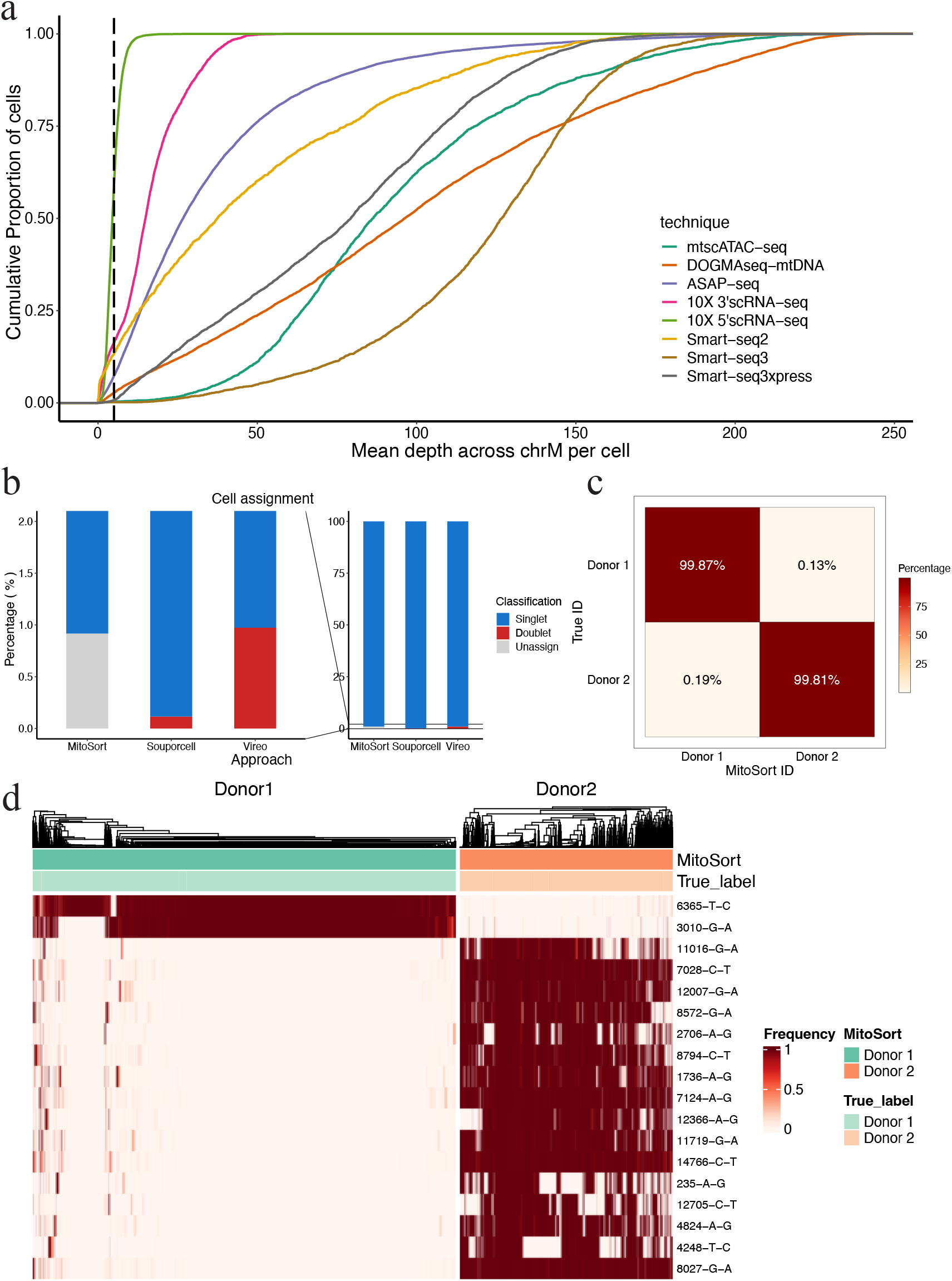
Extension of MitoSort to full-length scRNA-seq data. **a**, An Empirical Cumulative Distribution Function (ECDF) plot showing average sequencing depth across mitochondrial genome per cell for eight datasets using different sequencing techniques. **b,** Percentage of cell assignment of each tool on multiplexed Smart-seq3xpress dataset consisting of three donors. Both Souporcell and Vireo have false positive of doublet identification. **c,** Percentage of barcodes shared between dual indexes-based (rows) and MitoSort-based (columns) assignments in multiplexed Smart-seq3xpress data consisting of two donors. **d,** Heatmap showing the allele frequency of sample-specific mitochondrial germline variants (rows) across MitsoSort-assigned singlets (columns) in multiplexed Smart-seq3xpress dataset consisting of two donors. Top legend shows the donor assignments obtained from MitoSort and dual indexes.

**Supplementary Data Figure 5.**
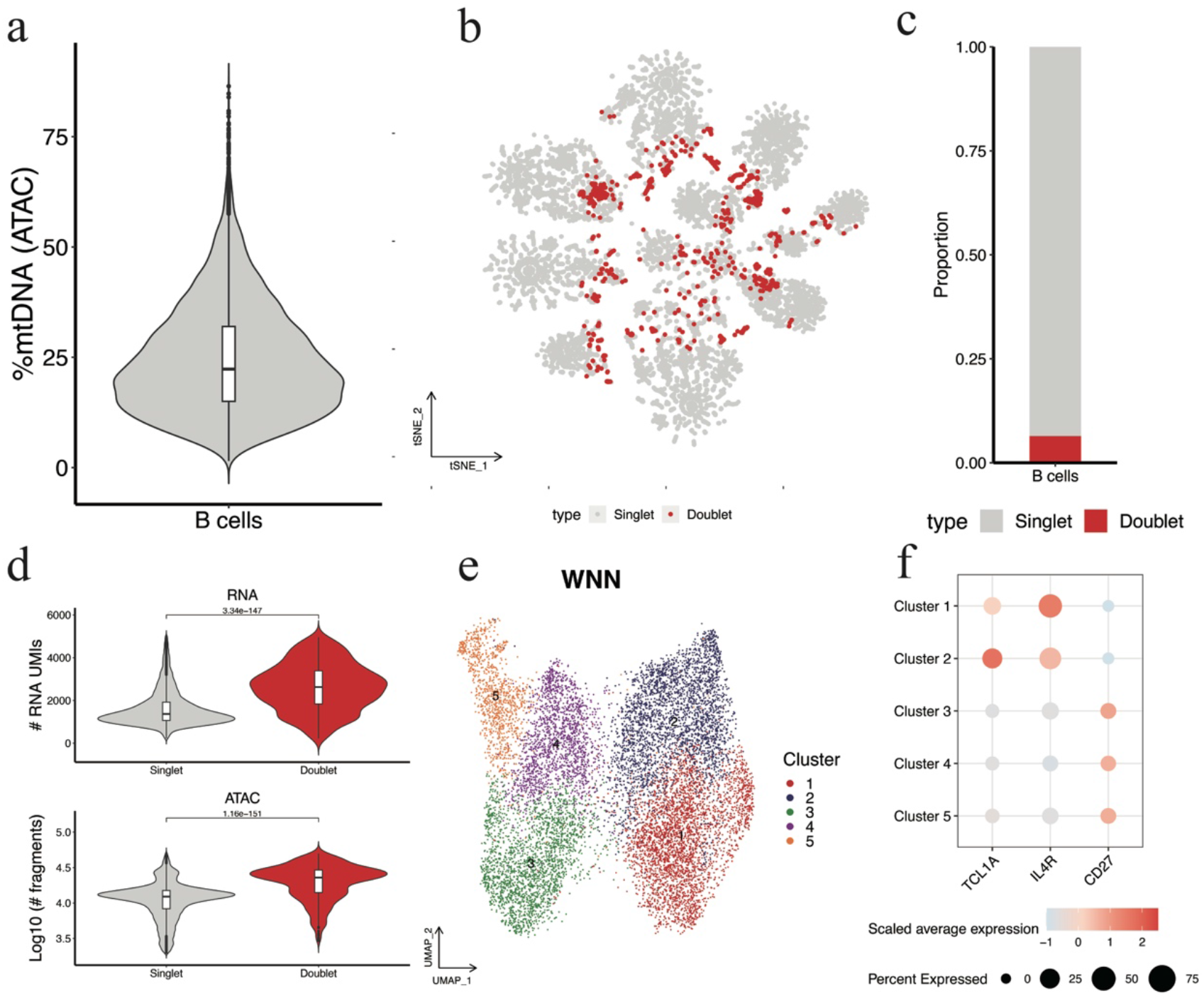
Analysis of multi-omics data of multiplexed B cells. **a**, Violin plot showing distribution of percentage of mtDNA reads per cell in the scATAC-seq library. **b,** tSNE plot of cell barcodes based on mitochondrial germline mutation profiles. Each dot represents a cell barcode and is colored based on the singlet/doublet classification by MitoSort. **c,** Bar plot showing proportion of cell barcodes that are classified as singlet and doublet by MitoSort. **d,** Violin plots showing the distribution of the number of RNA molecules (top) and ATAC fragments (bottom) per cell in cell barcodes classified as singlet and doublet. **e,** UMAP of weighted-nearest neighbor (WNN) graph for RNA and ATAC-seq modalities. Each dot represents a cell colored by its cluster. **f,** Dot plot showing expression of marker genes in each cluster. Dot size indicates the proportion of cells expressing the gene and dot color represents the scaled average expression level.

**Supplementary Table 1.** Data resource.

| DataSet | Library | Author |
| --- | --- | --- |
| GSE142745 | mtscATAC-seq | Lareau et al.2018 |
| GSM4732140 | mtscATAC-seq | Mimitou et al.2020 |
| GSE156474 | mtscATAC-seq | Mimitou et al.2020 |
| GSE156478 | mtscATAC-seq | Mimitou et al.2020 |
| GSE197037 | ASAP-seq | Rückert et al. 2022 |
| GSE200417 | DOGMA-seq | Xu et al. 2022 |
| GSE133549 | smart-seq2 | Mereu et al. 2020 |
| PRJCA016782 | RNA+ATAC multiome | Generated in this study |

